# arcasHLA: high resolution HLA typing from RNA seq

**DOI:** 10.1101/479824

**Authors:** Rose Orenbuch, Ioan Filip, Devon Comito, Jeffrey Shaman, Itsik Pe’er, Raul Rabadan

## Abstract

**Motivation:** Human leukocyte antigen (HLA) locus makes up the major compatibility complex (MHC) and plays a critical role in host response to disease, including cancers and autoimmune disorders. In the clinical setting, HLA typing is necessary for determining tissue compatibility. Recent improvements in the quality and accessibility of next-generation sequencing have made HLA typing from standard short-read data practical. However, this task remains challenging given the high level of polymorphism and homology between the HLA genes. HLA typing from RNA sequencing is further complicated by post-transcriptional splicing and bias due to amplification.

**Results:** Here, we present arcasHLA: a fast and accurate *in silico* tool that infers HLA genotypes from RNA sequencing data. Our tool outperforms established tools on the gold-standard benchmark dataset for HLA typing in terms of both accuracy and speed, with an accuracy rate of 100% at two field precision for MHC class I genes, and over 99.7% for MHC class II. Importantly, arcasHLA takes as its input pre-aligned BAM files, and outputs three-field resolution for all HLA genes in less than 2 minutes. Finally, we discuss evaluate the performance of our tool on a new biological dataset of 447 single-end total RNA samples from nasopharyngeal swabs, and establish the applicability of arcasHLA in metatranscriptome studies.

**Availability:** arcasHLA is available at https://github.com/RabadanLab/arcasHLA.

## 1 Introduction

Human leukocyte antigens encode the proteins that make up the major compatibility complex (MHC). MHC class I (HLA-A, B, and C) presents endogenous antigens on the surface of all nucleated, somatic cells to cytotoxic T-cells, triggering apoptosis if the protein is not recognized as self. MHC class II (including HLA-DPB1, DQB1, and DRB1), constitutively expressed by certain immune and epithelial cells, performs the same role but presents exogenous proteins to helper T-cells which mediate adaptive immune response [24].

HLA genes are the most polymorphic regions in the human genome with over 12,000 known alleles across 38 genes [30]. Pathogen-driven selection may explain this level HLA diversity: variation of residues in the binding region allows for a greater variety of peptides that can be bound and presented. Populations in areas with a wider variety of pathogens show increased HLA diversity [29] and heterozygous individuals show both greater resistance towards infectious agents and greater fitness than homozygotes [8; 34; 27].

With the advent of immunotherapy, HLA typing and expression level quantification is increasingly important for cancer research. Immunotherapy depends on the ability of the patient’s HLAs to effectively bind and present tumor neoantigens on the cell surface [9]. Following immunotherapy, clonal selection favors tumor cells with a loss of HLA heterozygosity (LOH) or silencing of the HLA loci. Although past methods look to copy number variations in whole exome sequencing to determine LOH [23], RNA sequencing may give a more accurate picture of HLA expression in tumor cells, particularly if HLA expression is altered as a result of interruptions in HLA regulatory pathways due to mutations or epigenetic modifications.

High resolution typing of HLA alleles is also imperative for the determination of tissue compatibility. HLA nomenclature (e.g. A*02:01:01:02L) consists of four successive fields: allele group, protein type, synonymous changes in coding regions, and changes in non-coding regions. The addition of a suffix denotes alterations in expression. “High-resolution” genotyping is used to determine an individual’s serotype, solving sequencing ambiguities in the peptide-binding region (exons 2 and 3 for class I and exon II for class II). Consequently, most sequenced HLA alleles are partial, missing exonic sequences outside of this region. However, differences in the peptide-binding sequence are not the only source of variation between alleles: expression levels, particularly that of HLA-C, are associated with allotype. Differences in expression can be exploited to salvage a match: if a donor has a single mismatched allele, there is decreased risk of transplant rejection if this allele is expressed at lower levels [28]. In short, accurate typing is necessary to determine compatibility between individuals.

Specialized methods of typing HLAs, including Sanger sequencing and PCR enrichment of the HLA loci, are expensive and time-consuming, given the sample size necessary for effective donor banks and association studies. Thus, methods using standard NGS reads with minimal loss of accuracy and resolution are useful. However, typing with short reads is made complicated by the high level of homology between both HLA genes and alleles, some of which differ by only a single base. In addition, there exist paralogous HLA pseudogenes, one of which has been shown to interfere with typing from genomic sequencing [17]. Some pseudogenes have detectable expression levels which interfere with RNA typing [22].

In the last few years, multiple tools that type HLAs from whole genome sequencing (WGS), whole exome sequencing (WES), and RNA sequencing have been published, with improving benchmark performance and resolution (see Table 1). These HLA typing tools attempt to find the one or two alleles that best explain the sampled reads, either by comparing assembled contigs or aligning reads directly to an HLA reference. Although a plethora of tools optimized for typing from WES and WGS fall into either category, most current tools for RNA sequencing, including seq2HLA[5], OptiType[33], PHLAT[3], are alignment-based. The latest RNA-dedicated HLA typer, HLAProfiler, takes a novel approach to graph-based alignment, breaking the HLA transcripts into k-mers and constructing a taxonomic tree used to filter reads [7]. To find an individual’s genotype, observed k-mers are compared to profiles built from simulated reads. Tools also differ in the construction of their HLA reference: some tools, such as seq2HLA and OptiType, limit their reference to peptide-binding exons and flanking regions while others use a combination of coding and genomic sequences. For the purposes of serotyping, changes outside of the peptide-binding region should be considered because alleles with the same peptide-binding sequence may have different protein types. In addition, limiting the number of exons considered increases the occurrence of ambiguous typing.

**Table 1.** Overview of tools cited in this paper.

| Tool | Input | MHC class | Resolution | Method | Partial | Input Format |
| --- | --- | --- | --- | --- | --- | --- |
| Polysolver | DNA | I | 4 fields | alignment | Y | BAM |
| HISAT-genotype | DNA | I and II | 4 fields | alignment followed by assembly | Y | FASTQ |
| xHLA | DNA | I and II | 4 fields | protein-level alignment | Y | BAM |
| OptiType | DNA/RNA | I | 2 fields | alignment | Y | FASTQ |
| PHLAT | DNA/RNA | I and II | 4 fields | alignment | N | FASTQ |
| seq2HLA | RNA | I and II | 2 fields | alignment | N | FASTQ |
| HLAProfiler | RNA | I and II | 4 fields | filtering followed by alignment | Y | FASTQ |
| **arcasHLA** | **RNA** | **I and II** | **3 fields** | **pseudoalignment** | **Y** | **BAM** |

arcasHLA takes an alignment-based approach, using both a coding DNA reference for complete alleles and a reference including all known alleles with all exon combinations of transcripts containing the peptide-binding region. This tool uses Kallisto [6], an RNA quantifier with a graph-based alignment feature, to assign reads to their compatible HLA transcripts. Allele abundance for each gene is quantified separately and the genotype that maximizes the number of reads aligned is selected from the most abundant alleles. Finally, homozygosity is determined using the ratio of minor to major nonshared read counts. As an optional step, partial alleles are typed in a similar fashion. Unlike other tools, population-specific allele frequencies are used as priors to distribute sampled reads within HLA compatibility classes in addition to breaking ties between ambiguous alleles (see Methods). arcasHLA outperforms other popular HLA RNA-sequencing typers such PHLAT, OptiType, seq2HLA, and HLAProfiler on paired-end benchmark samples (see Table 2).

**Table 2.** Concordance with gold-standard HLA typing of arcasHLA and other typers ran on 358 RNA-sequencing samples.

| Gene | % complete | OptiType | seq2HLA | PHLAT | HLAProfiler | **arcasHLA** |
| --- | --- | --- | --- | --- | --- | --- |
| A | 99.9% | 99.6% | 98.6% | 99.4% | 99.9% | **100.0%** |
| B | 99.9% | 99.4% | 94.8% | 93.4% | 99.0% | **100.0%** |
| C | 99.9% | **100.0%** | 95.1% | 94.3% | 99.6% | **100.0%** |
| DQB1 | 96.8% | - | 96.0% | 96.0% | **99.9%** | **99.9%** |
| DRB1 | 98.6% | - | 98.5% | 98.5% | 99.6% | **99.7%** |

## 2 Materials and methods

### 2.1 Database construction

#### 2.1.1 HLA reference

HLA and related sequences were obtained from the ImMunoGeneTics/HLA database, IMGT/HLA, compiled by the Immuno Polymorphism Database project [30]. These sequences include both classical and nonclassical MHC class I genes, MHC class II genes, HLA pseudogenes and some related non-HLA genes.

HLA nomenclature is divided into four fields: allele group, protein type, synonymous changes in coding regions, and changes in noncoding regions. Due to post-transcription splicing, changes in intronic regions cannot be confidently determined from mature messenger RNA. Excluding introns, we constructed databases of coding DNA for HLA alleles. Sequenced untranslated regions (UTRs), missing for many alleles, were included as noncontiguous sequences. Including these sequences in the reference identifies reads that map to both untranslated regions and coding sequences. Alleles with insertions or deletions causing a stop loss in the final exon were truncated if the sequence continuing into the UTR contained no changes. Many alleles from the same gene share much of their final exon and 3’ untranslated region, and reads containing this transition from coding to noncoding would only be attributed to the extended alleles. Thus, reads from the untranslated regions of the true alleles would be improperly assigned to these stop-loss alleles, interfering with typing.

A majority of the alleles archived in IMGT/HLA are not complete, missing exons, introns, and untranslated regions. Some HLA typing tools include partial alleles by extending the sequence with an allele’s nearest neighbor (Optitype) or looks at each exon individually. The method described here uses two separate references for typing complete and partial alleles. The former contains only transcripts for alleles with complete sequences, while the latter contains transcripts for all possible contiguous combinations of exons for all known alleles (e.g. 2-3, 1-2-3, etc).

#### 2.1.2 Allele frequencies

Two-field allele frequencies were retrieved from AlleleFrequencies Net Database (AFND) [13]. Only populations considered to be gold-standard, with allele frequencies that sum to 1 and a sample size ≥ 50, were used to build the database. These sample populations were grouped into broad population categories following the categorization laid out by The National Marrow Donor Program [15]. To account for alleles not seen in the selected population and those not reported on AFND, Dirichlet smoothing was applied to the allele frequencies, treating the entirety of the AFND data and IMGT/HLA database reference as priors.

### 2.2 Genotyping

#### 2.2.1 Read alignment

arcasHLA takes as input a mapped RNA-seq BAM file. After extracting chromosome 6 reads (and when applicable, extracting any additional reads aligned to HLA decoys, or chromosome 6 alternate sequences as well) from input, we perform a pseudoalignment of the extracted reads with Kallisto [6], a graph-based RNA-seq quantifier. Kallisto builds a de Bruijn graph from the reference transcriptome, in which each k-mer represents a k-length sequence and each edge adds an additional base, connecting the node to the next k-mer seen in the sequence. Each read is decomposed into k-length sequences and hashed into the reference index. The compatibility class of a given read is then defined as the set of reference transcripts that are compatible with every one of its constituent k-mers. This method avoids base-by-base alignment in favor of speed; thus the moniker “pseudoalignment.” Because Kallisto skips k-mers that provide no new information on the compatibility class of a read, it is less sensitive to sequencing errors if they happen to fall within any one of these redundant k-mers. Of note: this pseudoalignment method is also insensitive to novel alleles if the corresponding new variants lie along one of these conserved k-mer subsequences.

#### 2.2.2 Transcript quantification

Like most HLA typers, arcasHLA seeks to find the pair of alleles with maximal support among the observed reads originating from the HLA locus. Given the thousands of possible alleles for a single gene, pairwise comparisons, however, are computationally expensive and they fail to account for the similarity between different alleles. In order to narrow down the pool of possible alleles, arcasHLA exploits k-mer structures in transcript quantification, which, when combined with culling low-support allele transcripts, returns the allele pair (or possibly a single allele) that best explains the observed reads.

##### Division of counts

Traditionally, graph-based transcript quantifiers [26; 6] assign reads to equivalence classes of reference alleles, further sub-dividing reads within each compatibility class with equal weights among all the alleles in a given class. This approach may be beneficial when calculating differential expression of genes with many possible, equally-likely isoforms present in a single sample. To formalize, the setup for graph-based transcript quantifiers is as follows.

Let *A* be a set of reference alleles with lengths *l_a_* for *a* ∈ *A*, and *C* a set of observed compatibility classes consisting of subsets of *A*. For a given allele *i* ∈ *A*, we define *C_i_* ⊂ *C* as the set of compatibility classes which contain allele *i*. Each element *ω ∈ C_i_* is a compatibility class consisting of alleles in *A* with *i* ∈ *ω*. As such, the read count attributed to an allele *i* ∈ *A* with equal weights sub-division is then simply:

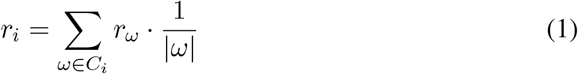

where |*ω*| denotes the number of alleles contained in the equivalence class *ω*, and *r_ω_* is the total count assigned to class *ω*.

arcasHLA performs genotyping calls with an iterative procedure that optimizes the read assignment to individual alleles. At the first step, our genotyping algorithm gives the option to distribute reads between alleles with weights proportional to population-specific allele frequencies. The largest benefit of this approach is narrowing the pool of possible alleles as well as breaking ties between alleles that are indistinguishable given the sampled reads. Given such priors *p* = (*p_i_*)_*i∈A*_, the count attributed to allele *i* is thus

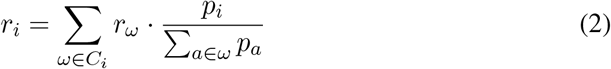

Subsequently, these counts are normalized by the allele length and converted into transcript abundances 0 ≤ *α_i_* ≤ 1 for each allele *i*:

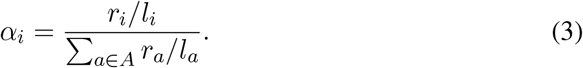

##### Maximizing the proportion of explained reads

As with Kallisto [6], the likelihood of a specific attribution of reads to alleles given by *α* = (*α_i_*)_*i∈A*_ is proportional to

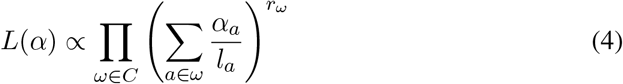

In order to find the allocation of reads to alleles that maximizes the likelihood function (Eq. 4), we follow an iterative procedure similar to Kallisto’s, with some essential differences.

First, we restrict the equivalence classes obtained from the reference de Bruijn graph construction gene by gene, and perform genotyping independently for each gene (namely, using our notation, we consider separately *A_HLA–A_, A_HLA–B_, A_HLA–C_*, …).

Second, instead of numerically solving for the maximum likelihood of (Eq. 4), we adopt a strong constrained approach consistent with our goal of outputting at most two alleles for each HLA gene. Reads in each class are iteratively reallocated based on abundances from the previous iteration, but after an empirically optimized 10 and 4 iterations for paired-end and single-end respectively, alleles with abundances lower than one tenth of the maximum observed abundance are dropped according to the following constraint:

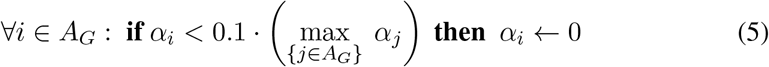

for each gene *G* in MHC class I and class II.

The 10 % threshold, previously determined by HISAT-genotype [19] for use with whole-genome sequencing, assumes that the abundance of the minor allele does not fall below a tenth of the major allele’s abundance. When applied to RNA sequencing, this allows for a large range in the natural variation between major and minor allele expression as well as differences in read counts due to sequencing and amplification.

The iterative read re-allocation in arcasHLA is as follows:

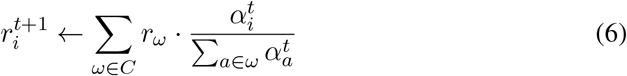

for all iterations *t* until convergence. Here, the upper indices denote the respective allele abundances or reads at the specified iteration. Next, these counts are normalized by transcript lengths and converted back into abundances:

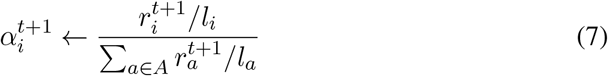

With each updated estimate, a higher proportion of reads are distributed to the alleles with the highest abundances and the lowest abundance alleles are culled, per (Eqn. 5).

Like HISAT-genotype [19], we use SQUAREM [35] to accelerate the convergence. The read allocation is considered to converge when the difference in abundance from the previous iteration to the current is below 10^−7^ with a maximum of 1000 iterations. Indeed, arcasHLA has been shown to always meet the convergence criterion in both of our test datasets, with the total number of iterations beneath 100 over all of runs. At the end of the arcasHLA iteration procedure, the remaining alleles are those that explain the highest proportion of reads aligned to a gene’s alleles.

#### 2.2.3 Selecting the most likely genotype

Ideally, after convergence and filtering out the low abundance alleles, a single allele is left for homozygous individuals and two alleles for the heterozygous ones. However, due to high levels of homology between certain alleles, particularly beyond the two field resolution, alleles may be indistinguishable given the observed reads and more than two likely alleles may be returned. In order to further narrow down the pool to exactly two alleles, the pair that explains the greatest proportion of reads is selected. Finally, we include a check for homozygosity by assessing the two alleles’ non-shared read counts. If the minor-to-major ratio of non-shared allele counts lies below an empirically-optimized threshold of 15%, the individual is predicted to be homozygous for the major allele. Otherwise, the individual is predicted to be heterozygous for the top ranking pair.

#### 2.2.4 Partial allele typing

Partial allele typing is included as an optional step. Extracted reads are aligned to the reference containing transcripts for complete and partial alleles. Counts are divided by gene and by included exons. Possible partial alleles are first identified by running transcript quantification on the peptide-binding exon transcripts. Next, arcasHLA iterates through the set of exon combinations represented in the returned partial alleles. If a partial allele has fewer than 10 reads more than the complete minor allele in that region, it is discarded as it cannot be confidently determined to be a valid allele. Next, all combinations of remaining partial alleles and the predicted complete alleles are considered. If a pair with one or more partial alleles explains a greater proportion of reads in any of these exon regions than the predicted complete genotype, it is returned as the most likely genotype. If more than one partial-containing pairs explains the same amount of reads, allele frequencies are used to break the tie.

### 2.3 Datasets

#### 2.3.1 Benchmark dataset: 1000 Genomes

HLA-A, B, C, DRB1, and DQB1 for 1,267 of the 1000 Genomes individuals were typed using Sanger sequencing based on the IMGT/HLA database from 2009 [14]. Only the peptide-binding region for each gene was sequenced. As previously stated, multiple alleles can share the same binding region sequence, and thus a list of equivalent alleles is reported. Since 2009, IMGT/HLA has expanded their database to more than four times as many alleles. Like HLAProfiler, we used the latest list of ambiguous alleles provided by IMGT/HLA to update the ground truth to reflect version 3.33.0.

mRNA sequencing for 358 of these samples is provided by the Geuvadis project, representing five of the 1000 Genomes populations (CEU, FIN, GBR, TSI, and YRI)[20]. These samples are generally high in quality with a mean RNA integrity number (RIN [31]) of 9.1 (ranging from 6.2 to 10), and a mean of 58.5*M* reads mapped to the hg19 reference (ranging from 17*M* to 163.5*M* reads). Reads are paired-end, and 75 base pairs (bp) in length. 25.1% and 14.8% of these individuals are homozygous for at least one gene at two fields in resolution for MHC class I and MHC class II respectively.

We ran arcasHLA on these samples IMGT/HLA v3.24.0, the version used by HLAProfiler [7]. This version was selected by Buchkovich, instead of the latest version at the time of HLAProfiler’s development v3.26.0, to increase the number of partial alleles in the dataset to demonstrate the tool’s ability to call partial alleles. In addition to updating the ground truth with allele ambiguities, calls were updated with the high-resolution typing using Ilumina TruSight provided by HLAProfiler. For comparisons with the 1000 Genomes dataset, we report the concordance of arcasHLA with the updated truth along with the rates of concordance of seq2HLA, OptiType, PHLAT, and HLAProfiler provided by Buchkovich (Tab. 2).

As a further test of arcasHLA’s accuracy, we downsampled all 358 samples to 5 million and 2.5 million reads. In addition, following PHLAT’s methodology, we treated the 1000 Genomes samples as single-end for both the original samples and the lower read count samples.

#### 2.3.2 New biological dataset: the Virome of Manhattan

We ran arcasHLA on a set of 447 single-end total RNA-sequencing samples collected from nasopharyngeal swabs from 69 healthy individuals enrolled as part of a DARPA-funded project entitled “The Virome of Manhattan: a Testbed for Radically Advancing Understanding and Forecast of Viral Respiratory Infections” [4; 12].

##### Sample collection and preparation

Nasopharyngeal samples were collected using minitip flock swabs and stored in tubes with 2 *ml* DNA/RNA Shield (Zymo Research, R1100-250) at 4-25 ° *C* for up to 30 days and then aliquoted into two 2 *ml* cryovials and stored at −80 °*C*. RNA was extracted from 200 *μl* of each stored sample using the Quick-RNA MicroPrep Kit (Zymo Research, Irvine, CA). Eluted RNA was then quantified and assessed for quality using Agilent Bioanalyzer (Santa Clara, CA), and the remaining quantity was sequenced with Illumina following the Ribo-Zero rRNA Removal Kit, target 30M single-end 100bp reads.

##### Sample processing

The individuals in the Virome study represent a heterogeneous cohort with self-reported and SNP-validated race/ethnicity (using the population clusters from the ExAC data set, [21]) from the African-American, Caucasian, Asian, Hispanic and Native American groups. As such, known population-specific allele frequency priors were passed to arcasHLA on this dataset executed in single-end mode from input BAM files mapped with STAR v.2.5.2b [11] to human reference GRCh37 [1]. In contrast to the high quality, homogeneous samples from the benchmark dataset, the Virome samples have a mean RNA integrity number (RIN [31]) of 7.0 (ranging from 1.0 to 9.9), and a mean of 22.2*M* reads mapped to the human GRCh37 reference (ranging from 5.9*M* to 68.2*M* reads).

##### Ground truth for typing comparison

We established the HLA genotyping ground truth for the Virome dataset using an assortment of *in silico* tools which attain high concordance with deep targeted sequencing validation protocols: xHLA [37], HISAT-genotype [19], OptiType [33] and Polysolver [32] – that we ran on whole exome sequencing (WES) data processed with the xGEN-Illumina platform (at 60 × 25*M* target PE 100bp reads) and extracted from saliva samples drawn independently from the nasopharyngeal swabs in our cohort.

Since we required both MHC class I and MHC class II predictions to test the full capability of arcasHLA, we resorted to setting xHLA’s two-field calls as the true Virome genotypes. We checked the concordance on the Virome WES data between xHLA and two other reliable tools: OptiType and Polysolver, which only return class I genes. On average, for HLA-A, -B, and -C genes, the two HLA calling methods showed good agreement with xHLA (97.9% for OpiType, 92.7% for Polysolver – see Table 3 for complete results). Further, in order to optimize speed and memory usage, we first used the HISAT-genotype `extract_reads` function (which builds on the HISAT aligner [18]) to extract reads mapping to the HLA locus before genotyping with xHLA. xHLA calls MHC class I HLA-A, -B, and -C and MHC class II HLA-DPB1, -DQB1, and -DRB1. HISAT-genotype and arcasHLA were run using IMGT/HLA databse v. 3.26.0. According to xHLA calls, the Virome individuals show lower rates of homozygosity than in the benchmark set with rates of homozygosity of 14.5% and 10.1% for MHC class I and MHC class II respectively.

**Table 3.**
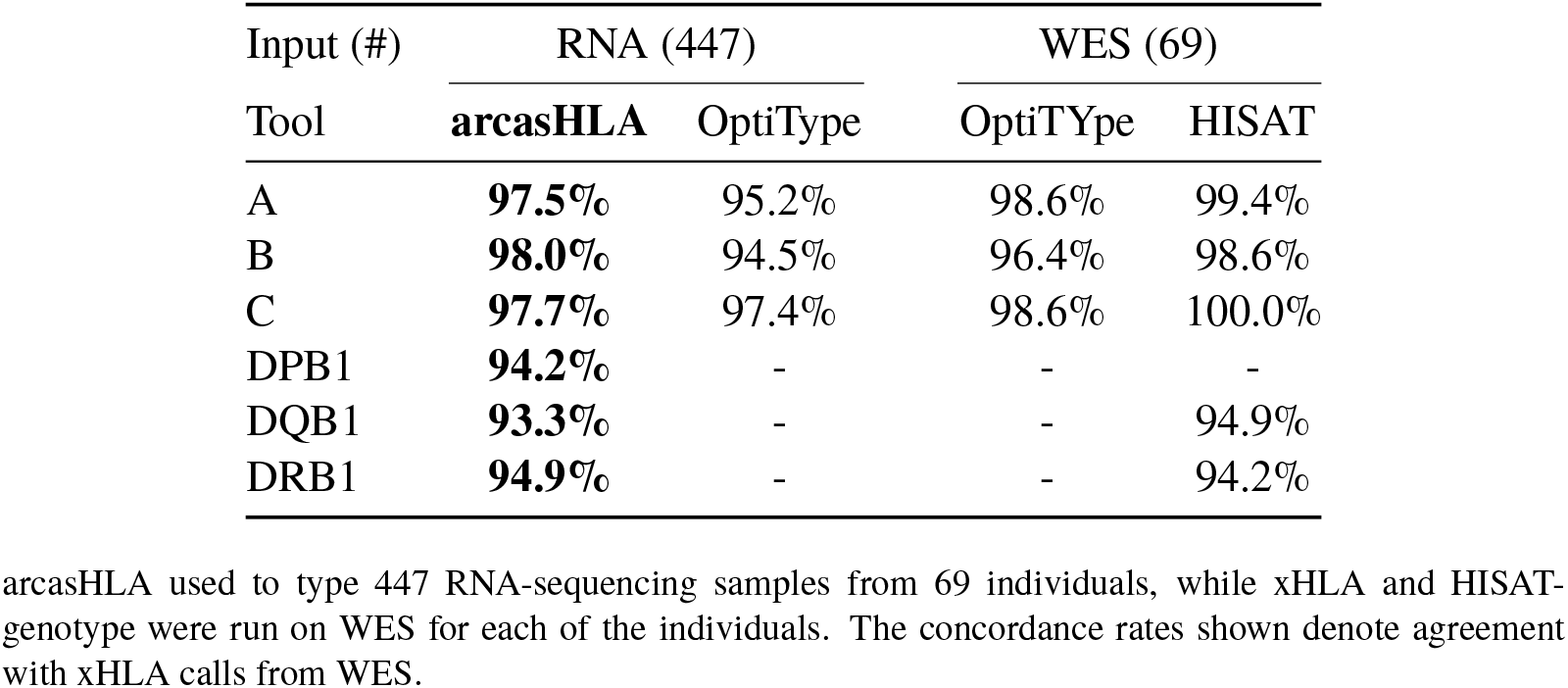
Concordance of calls from Virome samples with xHLA genotypes.

### 2.4 Implementation and availability

arcasHLA is as a command line tool written in Python available from the public GitHub repository https://github.com/RabadanLab/arcasHLA. This software is divided into four steps (Fig. 1). (1) Database construction takes fewer than 3 minutes on average and allows for the selection of a specific IMGT/HLA version given the commit hash. (2) Reads are extracted from previously sorted and indexed bam files. (3) Reads are aligned and allele abundances are quantified, followed by prediction of the most likely genotype. (4 optional) Reads are aligned to a reference containing partial alleles. Possible partial alleles are selected then compared with the genotype from the previous step.

**Figure 1.**
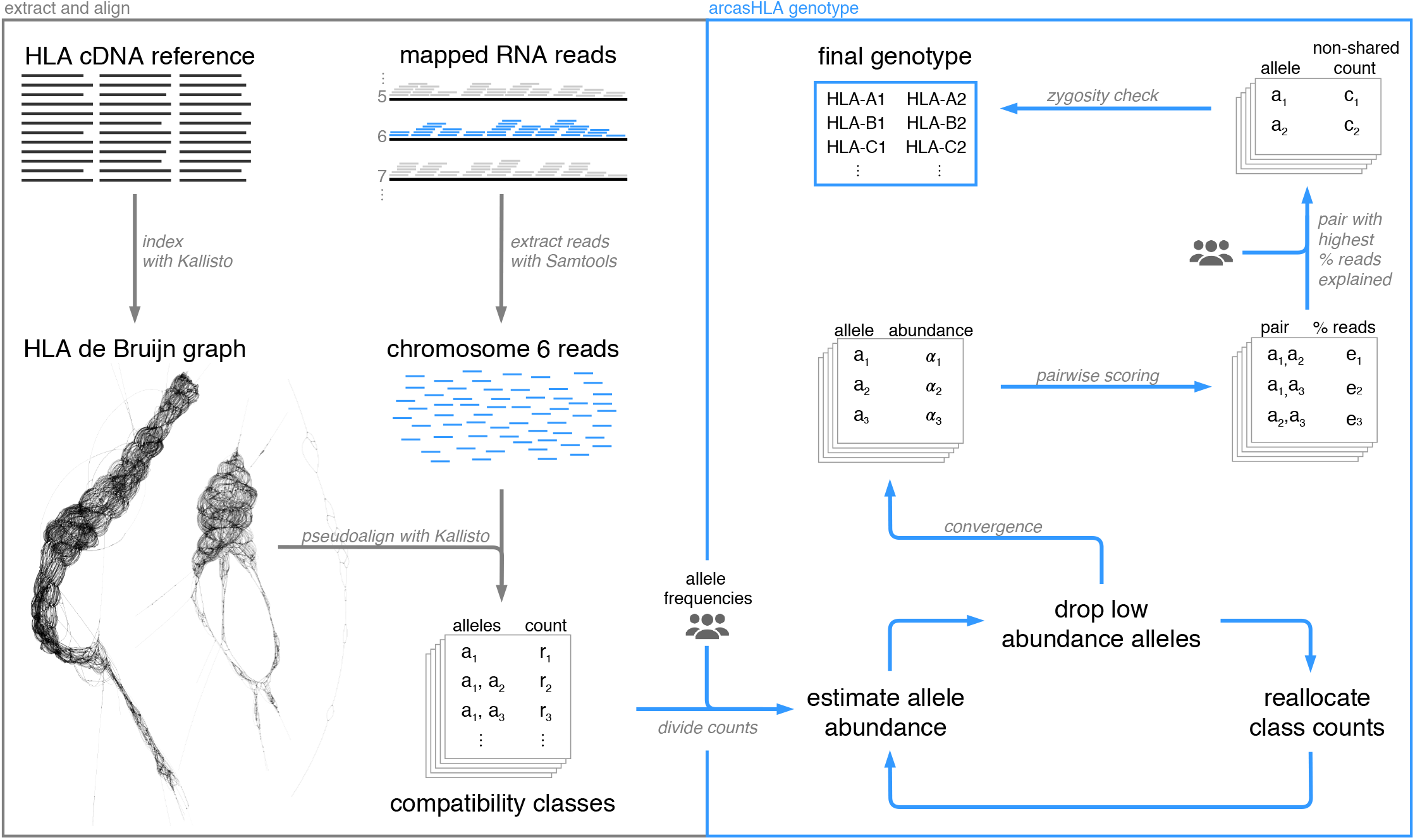
Overview of arcasHLA pipeline from alignment to genotyping. Visualization of HLA de Bruijn graph constructed using Velvet and Bandage.

The Geuvadis RNA sequencing of the 1000 Genomes individuals is available from ArrayExpress (E-GEUV-1). Pending consent from individuals enrolled in the Virome study (currently underway), the extracted reads mapping to chromosome 6 will be publicly released.

## 3 Results

### 3.1 Benchmark performance

When run on the 1000 Genomes benchmark set, arcasHLA achieves 100% accuracy for class I and above 99.7% accuracy for class II genes, outperforming other tools overall (Tab. 2). Errors are due to missing calls for two partial alleles, one DRB1 and one DQB1, incorrectly calling a complete allele for a single DRB1 allele. Overall, arcasHLA provides high levels of concordance for the HLA region using this benchmark set. The percent of complete alleles in the gold-standard set given the reference version 3.24.0 is provided in Table 1 as PHLAT and seq2HLA do not include partial alleles in their references. Consequently, their concordance rates are lower than the tools capable of partial allele typing.

#### 3.1.1 Runtime analyses

For computational analysis of arcasHLA, we randomly selected 30 samples from the 1000 Genomes benchmark dataset (Fig. 2). These samples, typed without the optional partial allele typing step, were analyzed on a Linux instance with 16 vCPUs and 64 GiB of memory using 8 threads per sample. All samples were genotyped in less than 2 minutes.

**Figure 2.**
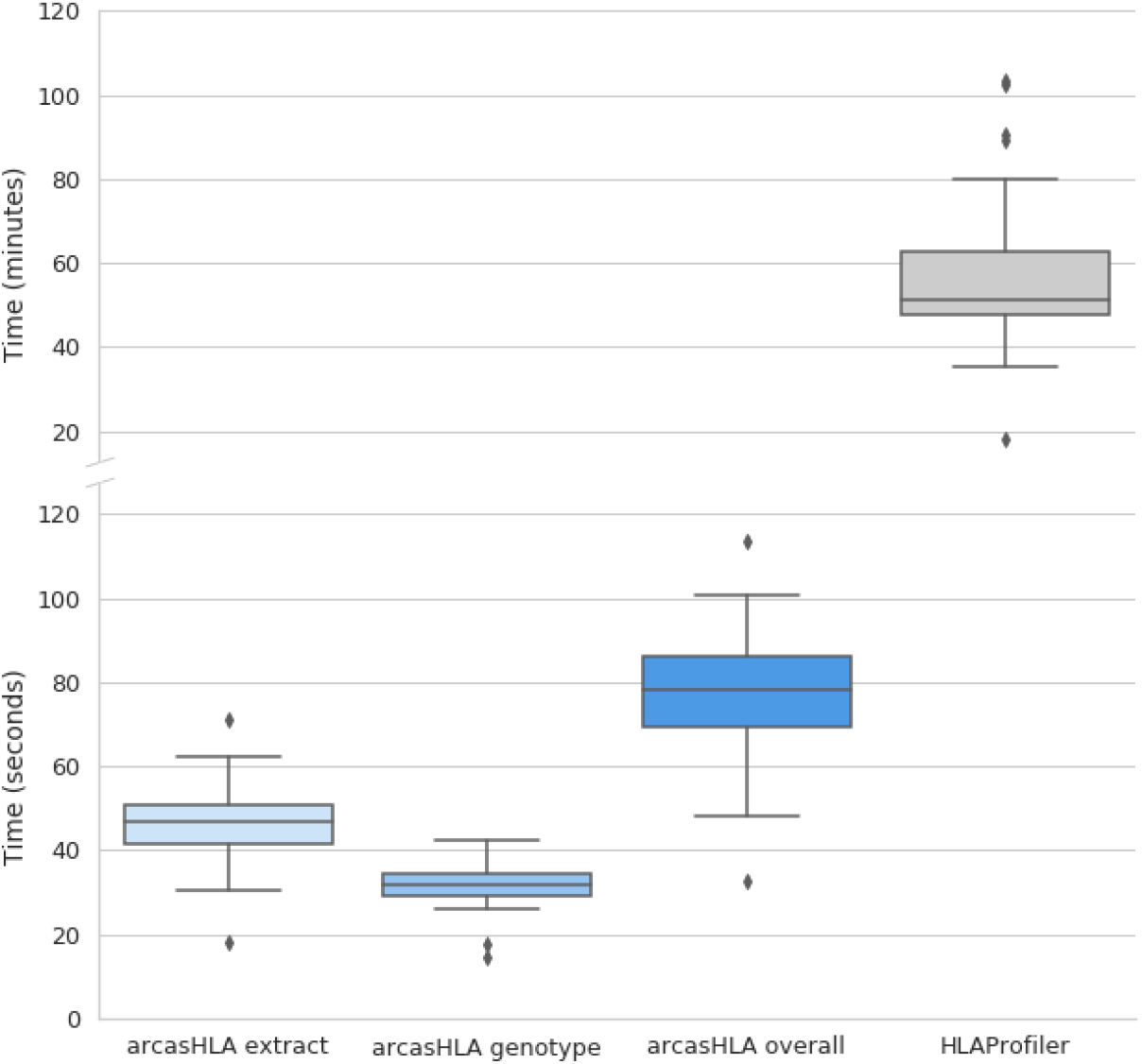
Runtime analysis on 30 randomly selected samples from 1000 Genomes dataset for arcasHLA (extract and genotype steps and overall runtime) and HLAProfiler.

HLAProfiler [7] is the top competitor for arcasHLA, as it is able to genotype both class I and class II MHC with high accuracy. arcasHLA, however, effectively achieves an order of magnitude runtime improvement over HLAProfiler when mapped RNA-seq reads are readily available, as HLAProfiler does not provide support for and does not benefit from pre-aligned sample input.

### 3.2 Performance on the Virome of Manhattan dataset

In spite of the lower quality metrics in the Virome dataset, arcasHLA yields high accuracy (Table 3): 94.8% for class I and 92.3% for class II.

Expression of MHC class II in the Virome samples can be attributed to the upper airway epithelial cells which are known to constitutively express MHC class II [36], and to the infiltration of leukocytes within the tissue lining the turbinates. Previous transcriptome analyses have shown that leukocyte markers are indeed expressed at low but detectable levels in samples from nasopharyngeal swabs [10].

Such specialized epithelial and immune cells are likely in the minority, however, which which may explain our tool’s lower accuracy result for class II. In fact, although arcasHLA was able to correctly predict MHC class II alleles for a majority of the samples, it failed to call HLA-DQB1 for several samples with an RIN of 1 and without any mapped reads to the DQB1 reference alleles.

We highlight the fact that the Virome samples were extracted from nasopharyngeal swabs and that they contain variable mixtures: human, bacterial and viral RNA (as detected by a BLAST search of the un-mapped reads [2]). RIN score is impacted by the proportion of human to prokaryotic reads, factoring in the 28S to 18S rRNA ratio. The variable sampling depth of the nasal cavity is another source of RIN variation and it can have a considerable impact, as mentioned above, on the read count and coverage of HLA genes. It is likely that another source of error stems from the fact that the protocol used in the Virome study was single-end sequencing (which is known to generate a less accurate mapping). Upon further analysis, arcasHLA occasionally misses calls because it fails to distinguish between very similar alleles which only differ in a few bases. Although this usually accounts for the variation of three field calls between typers, inability to resolve difference between ambiguous alleles can be exacerbated by single-end reads.

In spite of these study limitations, we report that HLA calling can still be successfully performed *in silico* from low-RIN samples with relatively low coverage of the HLA locus (Fig. 3).

**Figure 3.**
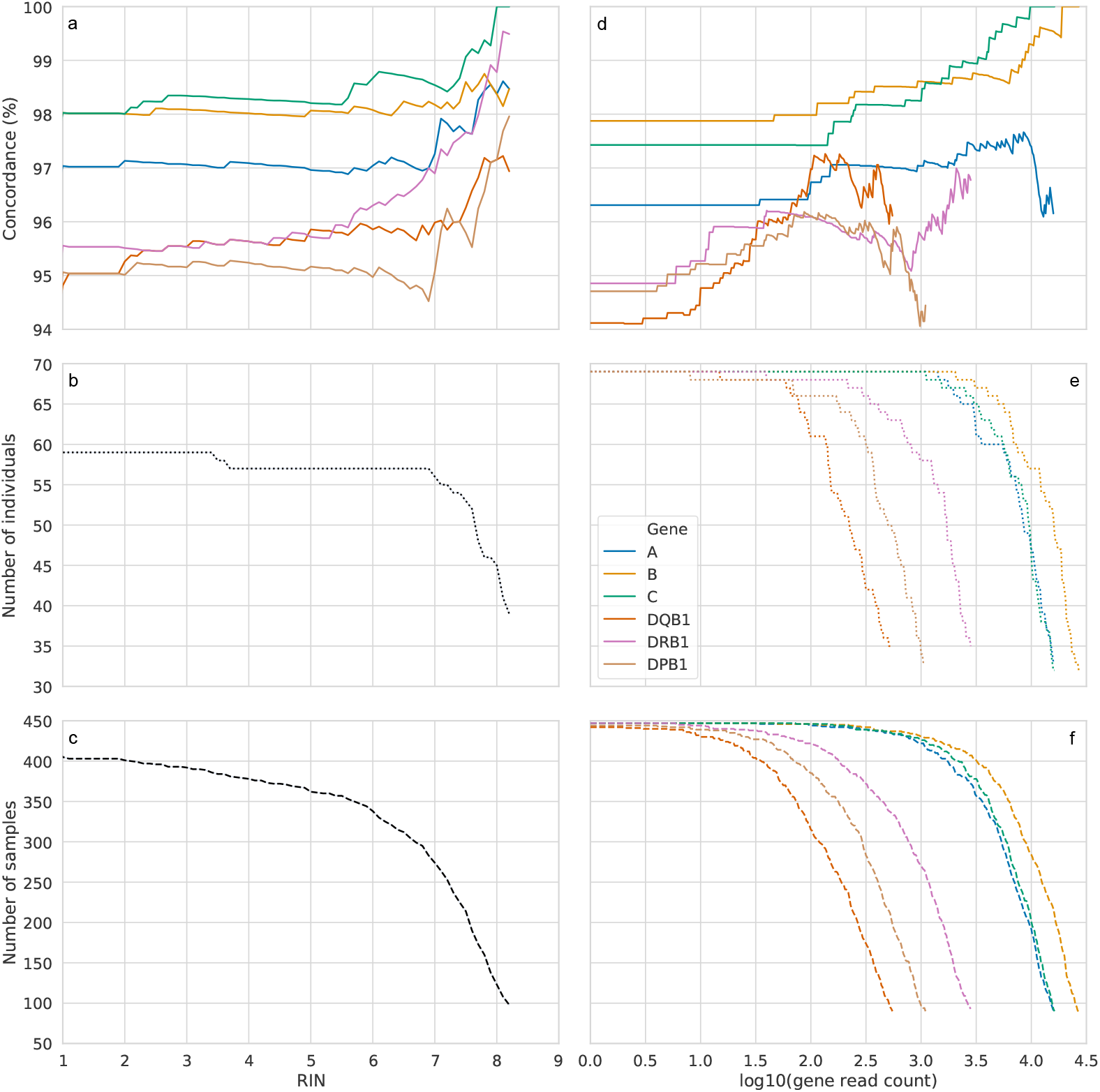
Accuracy rates restricted to Virome samples that meet the specified threshold criteria for (a) RIN and (d) log-normalized reads by HLA gene, truncated when the number of samples dropped below 89, approximately one fifth the total sample size. Panels (b) and (e) show the number of individuals with samples that meet the respective criteria and (c) and (f) show the number of samples remaining. The drops in accuracy rates for each gene at the highest read count threshold are explained by the one or two calling errors arcasHLA makes in less than 3 % of the samples.

## 4 Discussion

Accurate high-resolution HLA typing is imperative for determining tissue and hematopoietic compatibility. Typing from NGS reads helps narrow down the field of donors in a preliminary search, and is a boon to large-scale association studies where specialized assays prove too time-consuming and expensive. However, typing from shorter reads is made difficult by the high level of homology between HLA alleles and genes. Typers must be able to resolve ambiguities even at times when information is limited, such as when typing samples with low read count, short length reads, or single-end sequencing.

We have adapted transcript quantification algorithms to aid in typing of HLAs, a method which could be extended to type other highly polymorphic regions. arcasHLA performs at or near 100% accuracy on the gold-standard benchmark set, outperforming all other typers that run on RNA sequencing data. We have also validated our tool on a new biological dataset from a metatranscriptomic study of human nasopharyngeal swabs, showing how low read counts and low quality reads (as measured by the RNA integrity number) can affect the ability to type the MHC locus *in silico*.

A recent typer, HISAT-genotype, currently optimized for typing WGS, includes allele abundance quantification using methods from transcript quantifiers such as Kallisto. arcasHLA takes this approach further, adapting RNA-sequencing based transcript quantification to the HLA region. Taking advantage of Kallisto’s ultra-fast pseudoalignment, arcasHLA types HLA alleles both quickly and accurately. Like HISAT-genotype and the earliest release of Sailfish, arcasHLA uses a method for accelerating EM, SQUAREM. However, this method shows some level of instability in some rare cases. We plan to update the transcript quantification method with other EM accelerating algorithms in the future.

Although many typers use allele frequencies to break ties between ambiguous alleles, arcasHLA uses them to aid in the division of reads before quantification. The use of allele frequencies narrows down the pool of possible alleles, and lowers the impact of noisy reads and sequencing errors due to forgoing base-by-base alignment. Despite relying upon frequencies of alleles observed in particular populations, this method is still able to call both rare and partial alleles.

As benchmark performance for new tools approach 100% accuracy for standard benchmark tests, other criteria must be used to distinguish between them. While HLAProfiler, OptiType and arcasHLA perform similarly well, OptiType does not type class II genes and HLAProfiler does not accept single-end reads. In addition, arcasHLA has substain-tially faster runtimes on average. arcasHLA provides accurate, fast typing from RNA sequencing for both paired-end and single-end reads with easily parsed output.

The development of HLA typing tools from DNA and RNA sequencing is limited by the availability of gold-standard, benchmark data sets. These sets, used to both develop and test these tools, have only two field resolution typing as well as unresolved ambiguity between alleles beyond the peptide-binding region. The development of typers with accurate calling beyond the second field is hampered by the lack of public data sets with quality NGS samples and highest resolution typing.

In recent years, aided by landmark developments in 16S rRNA sequencing, whole-genome shotgun metagenomic sequencing and total RNA sequencing, a whole body of work has begun to map out the critical importance of our microbiome in systemic immunity, development, homeostasis, disease and patient responses to immunotherapy [16; 25; 38]. As in our project on the Virome of Manhattan, we expect that future meta-transcriptomic studies of the human host will rely on *in silico* methods in order to disentangle human from bacterial reads and maximally extract biological signal from lower quality and highly heterogeneous bulk samples. In this setting, HLA typing of the host, which depends on such signals, is of important clinical relevence. arcasHLA has been validated here for use with bulk total RNA samples containing eukaryotic and prokaryotic mixtures, showing high concordance for MHC class I and class II with the top HLA calling tools. Indeed, arcasHLA is impacted minimally by low read counts, low quality (as measured by RIN) and single-end sequencing protocol.

In the future, we plan on adding confidence for the most likely genotype calls as well as a more robust check for zygosity that takes expected levels of noise into account. Along these lines, we are working on using arcasHLA for testing the loss of heterozygosity in tumors and verifying mutations called from genomic sequencing. Because arcasHLA is based on RNA transcript quantifiers, it is natural to extend its functionality to allele specific quantification post genotyping – a feature in the works for the next version of our software. Expression-level data may enable us to detect loss of HLA expression or silencing as a possible mechanism of immune evasion.

## Author Contributions

R.O. and I.F. wrote the manuscript with input from all of the authors. R.O. conceived the idea for the tool, and, along with I.P. and I.F., developed the models used. R.O. designed and wrote the software presented. I.F. conducted the analysis of the Virome samples. R.R. and J.S. provided the biological samples from the Virome project. D.C. performed the laboratory work on the Virome samples. R.R. and I.P. supervised the project.

## Acknowledgement

We thank Andrew Chen and Benjamin Schweinhart for insightful comments and suggestions on an early version of the draft. Additionally, we thank Ruthie Birger, Marta Galanti, Erik Ladewig, Haruka Morita and Minhaz Ud-Dean for useful discussions. We would also like to thank Adithya Paramasivam for his help with testing the code.

## References

[1] Aken BL, Ayling S, Barrell D, et al, Searle SMJ. 2016. Database 2016. The Ensembl gene annotation system doi:10.1093/database/baw093.

[2] Altschul S, Gish W, Miller W, Myers E, Lipman D. 1990. Basic local alignment search tool. J. Mol. Biol. 215:403–410.

[3] Bai Y, Ni M, Cooper B, Wei Y, Fury W. 2014. Inference of high resolution HLA types using genome-wide RNA or DNA sequencing reads. BMC Genomics doi:10.1186/1471-2164-15-325.

[4] Birger R, Morita H, Comito D, Filip I, Galanti M, Lane B, Ligon C, Rosenbloom D, Shittu A, Ud-Dean M, Desalle R, Planet P, Shaman J. 2018. Asymptomatic shedding of respiratory virus among an ambulatory population across seasons. mSphere doi:10.1128/mSphere.00249-18.

[5] Boegel S, Löwer M, Schäfer M, Bukur T, de Graaf J, Boisguérin V, Türeci Ö, Diken M, Castle JC, Sahin U. 2012. HLA typing from RNA-Seq sequence reads. Genome Medicine doi:10.1186/gm403.

[6] Bray NL, Pimentel H, Melsted P, Pachter L. 2016. Near-optimal probabilistic RNA-seq quantification. Nature Biotechnology 34:525–527. doi:10.1038/nbt.3519.

[7] Buchkovich ML, Brown CC, Robasky K, Chai S, Westfall S, Vincent BG, Weimer ET, Powers JG. 2017. HLAProfiler utilizes k-mer profiles to improve HLA calling accuracy for rare and common alleles in RNA-seq data. Genome Medicine doi:10.1186/s13073-017-0473-6.

[8] Carrington M, Nelson GW, Martin MP, Kissner T, Vlahov D, Goedert JJ, Kaslow R, Buchbinder S, Hoots K, Brien SJO. 1999. HLA and HIV-1: Heterozygote Disadvantage 283:1748–1752.

[9] Chowell D, Morris LG, Grigg CM, Weber JK, Samstein RM, Makarov V, Kuo F, Kendall SM, Requena D, Riaz N, Greenbaum B, Carroll J, Garon E, Hyman DM, Zehir A, Solit D, Berger M, Zhou R, Rizvi NA, Chan TA. 2018. Patient HLA class I genotype influences cancer response to checkpoint blockade immunotherapy. Science doi:10.1126/science.aao4572.15334406.

[10] Chu CY, Qiu X, Wang L, Bhattacharya S, Lofthus G, Corbett A, Holden-Wiltse J, Grier A, Tesini B, Gill SR, Falsey AR, Caserta MT, Walsh EE, Mariani TJ. 2016. The Healthy Infant Nasal Transcriptome: A Benchmark Study. Scientific Reports 6:1–11. doi:10.1038/srep33994.

[11] Dobin A, Davis C, Schlesinger F, Drenkow J, Zaleski C, Jha S, Batut P, Chaisson M, Gingeras T. 2013. STAR: ultrafast universal RNA-seq aligner. Bioinformatics 29:15–21.

[12] Galanti M, Birger R, Ud-Dean SMM, Filip I, Morita H, Comito D, Anthony S, Freyer GA, Ibrahim S, Lane B, Ligon C, Rabadan R, Shittu A, Tagne E, Shaman J. 2018. Longitudinal active sampling for respiratory viral infections across age groups. In preparation.

[13] González-Galarza FF, Takeshita LY, Santos EJ, Kempson F, Maia MHT, Da Silva ALS, Teles E Silva AL, Ghattaoraya GS, Alfirevic A, Jones AR, Middleton D. 2015. Allele frequency net 2015 update: New features for HLA epitopes, KIR and disease and HLA adverse drug reaction associations. Nucleic Acids Research 43:D784–D788. doi:10.1093/nar/gku1166.

[14] Gourraud PA, Khankhanian P, Cereb N, Yang SY, Feolo M, Maiers M, Rioux JD, Hauser S, Oksenberg J. 2014. HLA diversity in the 1000 genomes dataset. PLoS ONE 9. doi:10.1371/journal.pone.0097282.

[15] Gragert L, Madbouly A, Freeman J, Maiers M. 2013. Six-locus high resolution HLA haplotype frequencies derived from mixed-resolution DNA typing for the entire US donor registry. Human Immunology 74:1313–1320. doi:10.1016/j.humimm.2013.06.025.

[16] Grice EA, Segre JA. 2012. The Human Microbiome: Our Second Genome. Ann. Rev. Genomics Hum. Genet. 13:151–170.

[17] Kawaguchi S, Higasa K, Shimizu M, Yamada R, Matsuda F. 2017. HLA-HD: An accurate HLA typing algorithm for next-generation sequencing data. Human Mutation 38:788–797. doi:10.1002/humu.23230.

[18] Kim D, Langmead B, Salzberg SL. 2015. HISAT: a fast spliced aligner with low memory requirements. Nature Methods 12:357–360.

[19] Kim D, Paggi JM, Salzberg S. 2018. HISAT-genotype: Next Generation Genomic Analysis Platform on a Personal Computer. bioRxiv: 266197 doi:10.1101/266197.

[20] Lappalainen T, Sammeth M, Friedländer M, AC’t Hoen P, Monlong J, Rivas M, Gonzalez-Porta M, Kurbatova N, Griebel T, Ferreira P, Barann M. 2017. Transcriptome and genome sequencing uncovers functional variation in humans. Cancer Epidemiol Biomarkers Prev. 2014 4:1264–1272. doi:10.1016/S2214-109X(16)30265-0.Cost-effectiveness.

[21] Lek M, Karczewski KJ, Minikel EV, et al, Exome Aggregation Consortium. 2016. Analysis of protein-coding genetic variation in 60,706 humans. Nature 536:285–291. doi:10.1038/nature19057.

[22] Lonsdale J, Thomas J, Salvatore M, Phillips R, Lo E, Shad S, Hasz R, Walters G, Garcia F, Young N, Foster B, Moser M, Karasik E, Gillard B, Ramsey K, Sullivan S, Bridge J, Magazine H, Syron J, Fleming J, Siminoff L, Traino H, Mosavel M, Barker L, Jewell S, Rohrer D, Maxim D, Filkins D, Harbach P, Cortadillo E, Berghuis B, Turner L, Hudson E, Feenstra K, Sobin L, Robb J, Branton P, Korzeniewski G, Shive C, Tabor D, Qi L, Groch K, Nampally S, Buia S, Zimmerman A, Smith A, Burges R, Robinson K, Valentino K, Bradbury D, Cosentino M, Diaz-Mayoral N, Kennedy M, Engel T, Williams P, Erickson K, Ardlie K, Winckler W, Getz G, DeLuca D, Daniel MacArthur, Kellis M, Thomson A, Young T, Gelfand E, Donovan M, Meng Y, Grant G, Mash D, Marcus Y, Basile M, Liu J, Zhu J, Tu Z, Cox NJ, Nicolae DL, Gamazon ER, Im HK, Konkashbaev A, Pritchard J, Stevens M, Flutre T, Wen X, Dermitzakis ET, Lappalainen T, Guigo R, Monlong J, Sammeth M, Koller D, Battle A, Mostafavi S, McCarthy M, Rivas M, Maller J, Rusyn I, Nobel A, Wright F, Shabalin A, Feolo M, Sharopova N, Sturcke A, Paschal J, Anderson JM, Wilder EL, Derr LK, Green ED, Struewing JP, Temple G, Volpi S, Boyer JT, Thomson EJ, Guyer MS, Ng C, Abdallah A, Colantuoni D, Insel TR, Koester SE, A Roger Little, Bender PK, Lehner T, Yao Y, Compton CC, Vaught JB, Sawyer S, Lockhart NC, Demchok J, Moore HF. 2013. The Genotype-Tissue Expression (GTEx) project. Nature Genetics 45:580–585. doi:10.1038/ng.2653.NIHMS150003.

[23] McGranahan N, Rosenthal R, Hiley CT, Rowan AJ, Watkins TB, Wilson GA, Birkbak NJ, Veeriah S, Van Loo P, Herrero J, Swanton C, Swanton C, Jamal-Hanjani M, Veeriah S, Shafi S, Czyzewska-Khan J, Johnson D, Laycock J, Bosshard-Carter L, Rosenthal R, Gorman P, Hynds RE, Wilson G, Birkbak NJ, Watkins TB, Horswell S, Mitter R, Escudero M, Stewart A, Van Loo P, Rowan A, Xu H, Turajlic S, Hiley C, Abbosh C, Goldman J, Stone RK, Denner T, Matthews N, Elgar G, Ward S, Costa M, Begum S, Phillimore B, Chambers T, Nye E, Graca S, Al Bakir M, Joshi K, Furness A, Ben Aissa A, Wong YNS, Georgiou A, Quezada S, Hartley JA, Lowe HL, Herrero J, Lawrence D, Hayward M, Panagiotopoulos N, Kolvekar S, Falzon M, Borg E, Marafioti T, Simeon C, Hector G, Smith A, Aranda M, Novelli M, Oukrif D, Janes SM, Thakrar R, Forster M, Ahmad T, Lee SM, Papadatos-Pastos D, Carnell D, Mendes R, George J, Navani N, Ahmed A, Taylor M, Choudhary J, Summers Y, Califano R, Taylor P, Shah R, Krysiak P, Rammohan K, Fontaine E, Booton R, Evison M, Crosbie P, Moss S, Idries F, Joseph L, Bishop P, Chaturved A, Quinn AM, Doran H, Leek A, Harrison P, Moore K, Waddington R, Novasio J, Blackhall F, Rogan J, Smith E, Dive C, Tugwood J, Brady G, Rothwell DG, Chemi F, Pierce J, Gulati S, Naidu B, Langman G, Trotter S, Bellamy M, Bancroft H, Kerr A, Kadiri S, Webb J, Middleton G, Djearaman M, Fennell D, Shaw JA, Le Quesne J, Moore D, Nakas A, Rathinam S, Monteiro W, Marshall H, Nelson L, Bennett J, Riley J, Primrose L, Martinson L, Anand G, Khan S, Amadi A, Nicolson M, Kerr K, Palmer S, Remmen H, Miller J, Buchan K, Chetty M, Gomersall L, Lester J, Edwards A, Morgan F, Adams H, Davies H, Kornaszewska M, Attanoos R, Lock S, Verjee A, MacKenzie M, Wilcox M, Bell H, Hackshaw A, Ngai Y, Smith S, Gower N, Ottensmeier C, Chee S, Johnson B, Alzetani A, Shaw E, Lim E, De Sousa P, Barbosa MT, Bowman A, Jordan S, Rice A, Raubenheimer H, Proli C, Cufari ME, Ronquillo JC, Kwayie A, Bhayani H, Hamilton M, Bakar Y, Mensah N, Ambrose L, Devaraj A, Buderi S, Finch J, Azcarate L, Chavan H, Green S, Mashinga H, Nicholson AG, Lau K, Sheaff M, Schmid P, Conibear J, Ezhil V, Ismail B, Irvin-sellers M, Prakash V, Russell P, Light T, Horey T, Danson S, Bury J, Edwards J, Hill J, Matthews S, Kitsanta Y, Suvarna K, Fisher P, Keerio AD, Shackcloth M, Gosney J, Postmus P, Feeney S, Asante-Siaw J, Aerts HJ, Dentro S, Dessimoz C. 2017. Allele-Specific HLA Loss and Immune Escape in Lung Cancer Evolution. Cell 171:1259–1271.e11. doi:10.1016/j.cell.2017.10.001.

[24] Meyer D, Thomson G. 2001. How selection shapes variation of the human major histocompatibility complex: a review. Ann Hum Genet 65:1–26. doi:10.1046/j.1469-1809.2001.6510001.x.

[25] Obata Y, Pachnis V. 2016. The Effect of Microbiota and the Immune System on the Development and Organization of the Enteric Nervous System. Gastroenterology 151:836–844. doi:10.1053/j.gastro.2016.07.044.

[26] Patro R, Mount SM, Kingsford C. 2014. Seq Reads Using Lightweight Algorithms. Nature biotechnology 32:462–464. doi:10.1038/nbt.2862.Sailfish.

[27] Penn DJ, Damjanovich K, Potts WK. 2002. MHC heterozygosity confers a selective advantage against multiple-strain infections. Technical report.

[28] Petersdorf EW, Gooley TA, Malkki M, Bacigalupo AP, Cesbron A, Du Toit E, Ehninger G, Egeland T, Fischer GF, Gervais T, Haagenson MD, Horowitz MM, Hsu K, Jindra P, Madrigal A, Oudshoorn M, Ringdén O, Schroeder ML, Spellman SR, Tiercy JM, Velardi A, Witt CS, O’Huigin C, Apps R, Carrington M. 2014. HLA-C expression levels define permissible mismatches in hematopoietic cell transplantation. Blood doi:10.1182/blood-2014-09-599969.

[29] Prugnolle F, Manica A, Charpentier M, Guégan JF, Guernier V, Balloux F. 2005. Pathogen-driven selection and worldwide HLA class I diversity. Current Biology doi:10.1016/j.cub.2005.04.050.

[30] Robinson J, Halliwell JA, Hayhurst JD, Flicek P, Parham P, Marsh SG. 2015. The IPD and IMGT/HLA database: Allele variant databases. Nucleic Acids Research 43:D423–D431. doi:10.1093/nar/gku1161.

[31] Schroeder A, Mueller O, Stocker S, Salowsky R, Leiber M, Gassmann M, Lightfoot S, Menzel W, Granzow M, Ragg T. 2006. The RIN: an RNA integrity number for assigning integrity values to RNA measurements. BMC Molecular Biology 7. doi:10.1186/1471-2199-7-3.

[32] Shukla SA, Rooney MS, Rajasagi M, et al. 2015. Comprehensive analysis of cancer-associated somatic mutations in class I HLA genes. Nature Biotechnology 33:1152–1158. doi:10.1038/nbt.3344.

[33] Szolek A, Schubert B, Mohr C, Sturm M, Feldhahn M, Kohlbacher O. 2014. OptiType: Precision HLA typing from next-generation sequencing data. Bioinformatics doi:10.1093/bioinformatics/btu548. arXiv:1011.1669v3.

[34] Thursz MR, Thomas HC, Greenwood BM, Hill AV. 1997. Heterozygote advantage for HLA class-II type in hepatitis B virus infection. doi:10.1038/ng0997-11. ng0997-11.

[35] Varadhan R, Roland C. 2008. Simple and globally convergent methods for accelerating the convergence of any em algorithm. Scandinavian Journal of Statistics 35:335–353. doi:10.1111/j.1467-9469.2007.00585.x.

[36] Wosen JE, Mukhopadhyay D, Macaubas C, Mellins ED. 2018. Epithelial MHC Class II Expression and Its Role in Antigen Presentation in the Gastrointestinal and Respiratory Tracts. Front. Immunol. 9. doi:10.3389/fimmu.2018.02144.

[37] Xie C, Yeo Z, Wong M, et al, Venter JC. 2017. Fast and accurate HLA typing from short-read next-generation sequence data with xHLA. PNAS 114:8059–8064.

[38] Zitvogel L, Y M, D R, Kroemer G, F GT. 2018. The microbiome in cancer immunotherapy: Diagnostic tools and therapeutic strategies. Science 359:1366–1370. doi:10.1126/science.aar6918.

